# MoDPA: Inferring Modification-dependent Protein Associations from Uniformly Reprocessed Mass Spectrometry

**DOI:** 10.64898/2026.01.20.700550

**Authors:** Enrico Massignani, Natalia Tichshenko, Kevin Velghe, Lennart Martens

## Abstract

**Motivation:** Post-translational modifications (PTMs) are key regulators of protein function and cellular processes; however, the overall principles of PTM co-regulation and crosstalk remain to be fully understood. A major challenge in large-scale PTM crosstalk studies is the scarcity of data, which hampers the ability to reproducibly detect proteome-wide associations between modification sites.

**Results:** We present a new computational framework, named Modification-Dependent Protein Associations (MoDPA). To overcome the extreme sparsity and heterogeneity of PTM calls across experiments, MoDPA utilizes a variational autoencoder (VAE) to embed per-site detection profiles into a low-dimensional latent space that preserves covariation while denoising missing data. A PTM association network is constructed by correlating latent representations across experiments. Benchmarking against pulsed SILAC data shows that MoDPA can correctly capture the correlation between heavy- and light-labelled peptides. We apply MoDPA to a large sample of reprocessed public datasets and identify clusters of modified proteins involved in distinct biological pathways, suggesting that some modifications may preferentially regulate specific biological processes, but not others. In particular, we find clusters enriched in lysine acetylation, lysine methylation, and arginine deamidation (citrullination) sites, which contain proteins involved in protein synthesis, neuron development, and cellular senescence.

**Availability and Implementation:** All code is freely available on GitHub (https://github.com/CompOmics/MoDPAv1.0) and Zenodo (10.5281/zenodo.18310674). The resulting MoDPA-derived PTM network is available via the TabloidProteome website at https://iomics.ugent.be/tabloidproteome.

## Introduction

Proteins are involved in a wide range of biological functions, making them essential for the survival of any organism. However, proteins do not operate in isolation; they interact with one another in various ways. Protein-protein interactions (PPIs), which can be either direct (such as the formation of complexes) or indirect (such as participation in the same signaling cascade), alongside post-translational modifications (PTMs), are the primary mechanisms by which cells regulate protein functions and respond to stimuli [1–5].

Despite extensive research into protein associations, the interplay between different PTMs and how they influence each other remains a significant and as-yet unresolved area of inquiry [6]. To address this knowledge gap, we propose Modification-Dependent Protein Associations (MoDPA), a computational framework that infers PTM-PTM associations from the large-scale reprocessing of public mass spectrometry-based proteomics data.

Several studies have demonstrated that the phenomenon of co-occurrence across different experiments can be utilized to determine relationships between entities. For instance, consistent co-expression of genes or proteins across several datasets is routinely used in gene and protein co-expression analysis to infer PPIs and gene regulatory networks [7–11].

One limitation of proteomics compared to genomics is the high sparsity of the data, especially regarding PTM data [12]. As PTMs are sub-stoichiometric and often unstable or labile, modified peptides are particularly difficult to reproducibly identify across MS runs [13]. Furthermore, each protein has several possible PTM sites, resulting in matrices that have very high dimensionality and sparsity.

Nevertheless, Variational Autoencoder (VAE) models have been shown to be effective in reducing the dimensionality and sparsity of proteomics data, producing a low-dimensional latent space that reliably “summarizes” the original data [14,15]. This approach was successfully implemented in the FAVA approach to identify PPIs from a large, uniformly reprocessed set of MS data [16].

The approach we present here goes substantially beyond the original FAVA framework, in that MoDPA utilizes a VAE to encode post-translational modifications that are identified and quantified across various datasets into a latent space. For each pair of PTMs, the signed distance correlation between them is then assessed [17]. Ultimately, a weighted network is created where each node represents a modified site, and the signed distance correlations are encoded as the weights of the edges. The result is the first-ever, proteome-wide, modification-dependent protein association network.

We validated our approach by using Pulsed SILAC (pSILAC) data, whereby heavy and light labels are expected to change over time following a predictable trend: i.e., all heavy peptides are expected to increase, and all light peptides are expected to decrease as cells incorporate heavy isotope-labelled amino acids into proteins [18]. MoDPA correctly predicted positive associations within the two label groups (heavy and light), and negative associations across the groups.

Subsequent analysis of a large amount of real PTM data, uniformly reprocessed from public data sets, identified clusters of co-occurring PTMs that we then mapped to Reactome biological pathways [19], revealing both known and novel associations between specific PTMs and cellular processes.

## Methods

### Data reprocessing

MS raw data files were selected and downloaded from PRIDE [20] (Table S1). A dataset was considered amenable for reprocessing if: (i) it only included human samples, (ii) the fragmentation method was HCD, and (iii) the acquisition method was DDA.

Raw files were converted to.mgf format with ThermoRawFileParser [21] and processed with ionbot [22]. Enzyme specificity was set to trypsin, a maximum of two allowed missed cleavages, and mass tolerances for both precursor and fragment ions were set to 20ppm. The open modification search was performed in two steps: raw files were first searched with carbamidomethyl of Cysteine and oxidation of Methionine as predefined variable modifications, allowing up to one unexpected modification in addition. From these preliminary results, a list of the most abundant modifications in each LC-MS run was compiled. A second round of searches was then performed where a tailored list of predefined variable PTMs was provided for each LC-MS run, based on the 5 most abundant PTMs for that run. This procedure allowed us to identify peptides with a larger variety of PTMs. Fixed modifications were never specified. Results were filtered at PSM level for q-value < 0.01 (FDR < 1%).

### Quantification of PTMs

PTM quantification was performed through relative PSM counts. We defined this metric as the total PSM counts of all peptidoforms carrying a given PTM (i.e., absolute PSM counts) divided by the total PSM counts of all peptidoforms spanning that PTM site (regardless of modification). This ratio was used as a proxy for the stoichiometry of each PTM in the analysed MS runs. Peptidoforms with a PSM count of one were excluded from the calculations, as were PTMs with a total absolute count less than three.

The following modifications were extracted from the overall dataset to be analyzed: lysine acetylation (Unimod ID: 1); serine/threonine/tyrosine phosphorylation (Unimod ID: 21); serine/threonine/tyrosine dehydration (Unimod ID: 23); lysine/aspartic acid/glutamic acid carboxylation (Unimod ID: 299); lysine/arginine monomethylation (Unimod ID: 34); lysine/arginine dimethylation (Unimod ID: 36); lysine trimethylation (Unimod ID: 37); lysine ubiquitination (Unimod ID: 535); cysteine/histidine/lysine hydroxynonenal (Unimod ID: 53); lysine succninylation (Unimod ID: 64); arginine deamidation/citrullination (Unimod ID: 7).

The final co-expression matrix was filtered to include only PTMs observed in 25 or more experiments and only experiments with 25 or more identified PTMs.

### Variational Autoencoder (VAE) implementation

A VAE was used to compress the high-dimensional PTM abundance matrix into a lower-dimensional latent space.

In VAEs, the latent representations are regularized by minimizing the regularization loss, calculated as the Kullback-Leibler (KL) divergence between the latent and a prior, in this case a normal distribution with mean=0 and standard deviation=1. Reconstruction loss was computed as the cosine loss + 1 to ensure positive values (bounded by [0, 2]). The total VAE loss is composed of the sum of reconstruction and regularization losses.

Regarding the VAE architecture, the encoder consists of two dense layers using Rectified Linear Unit (ReLU) activation function, a dropout layer between the two hidden layers (dropout rate = 0.25), and finally two encoding layers for the mean (“enc_mu”) and for the log-transformed variance (“enc_logvar”), both using linear activation. The decoder consists of two dense layers using ReLU activation and an output layer using sigmoid activation, as the input consists of relative abundances, which are naturally bounded by [0, 1].

The VAE model was implemented in Keras [23], and trained using the Adam optimizer with a learning rate of 10^−3^. Sixteen models were trained in total and ranked based on the cosine similarity between the original and reconstructed data. The best-performing model, “compassionate_buck”, had hidden layers of size 4096 and 512, respectively, and a latent space of size 96 (Table S2).

### PTM Association inference

We compute all pairwise signed distance correlations (SDCs) and associated p-values between post-translational modifications (PTMs) within the VAE-generated latent space, using the *dcor* package. For each PTM pair, the SDC is obtained by copying the sign of the Pearson correlation coefficient to the distance correlation. This analysis yields a ranked list of PTM pairs, with higher SDC reflecting closer proximity in the latent space and, therefore, more similar abundance patterns. P-values were corrected for multiple testing using the Benjamini-Hochberg method.

### Analysis of Pulsed SILAC data for validation

Raw data files from pSILAC-labelled samples were downloaded from PRIDE (PXD021203, PXD037569, PXD042903) and searched with ionbot against the *Homo sapiens* canonical reference proteome (UniProtKB 2023_01). Cystein carbamidomethylation was indicated as a fixed modification. Methionine oxidation, N-terminal acetylation, ^13^C_6_-^15^N_4_ Arginine (R10), and ^13^C_6_-^15^N_2_ Lysine (K8) were included as variable modifications.

For the purpose of this analysis, R10 and K8 were treated as variable modifications rather than SILAC labels. The occupancy of R10 and K8 was measured over two time points.

PTM associations were then computed from the pSILAC PTM abundance matrix, as described above. Validation proceeded by rule-based edge labelling: (i) positive associations between two heavy-labelled residues or two light-labelled residues were marked valid; (ii) negative associations between one heavy-labelled and one light-labelled residue were marked valid; and (iii) all other associations were marked invalid. We then compared the enrichment of valid edges in the MoDPA network, a degree-preserving randomized network, and a fully randomized network. The network was randomized through an in-house Python script using the *NetworkX* library [24].

### Clustering and pathway enrichment analysis

Clustering was performed in Cytoscape with the Leiden algorithm (objective_function = modularity; resolution_parameter = 0.5; beta = 0.01; n_iterations = 2), using the SDCs values as weights [25,26]. The pathways enrichment analysis was performed using the Reactome Python client *reactome2py*. Of the clusters identified, Cluster 18 was not significantly enriched in any pathways, and Cluster 23 only contained 11 proteins; thus, these were excluded from the results.

## Results

### Pipeline description and validation

MoDPA utilizes PTM occupancy matrices as inputs to train a VAE [15]. PTM occupancy matrices are obtained by reprocessing publicly available MS data with ionbot, a machine learning-based peptide search engine (Fig. 1A). The VAE is then used to learn the distribution of the input data and create a meaningful low-dimensional representation (i.e., the latent space) (Fig. 1B). All-against-all correlations are calculated in the latent space to quantify similarity in abundance patterns between any two PTMs. In our implementation, we utilized SDC as the correlation metric, allowing us to detect nonlinear relationships between PTMs [17]. The resulting adjacency matrix is directly translated to a weighted functional association network (Fig. 1C).

**Figure 1.**
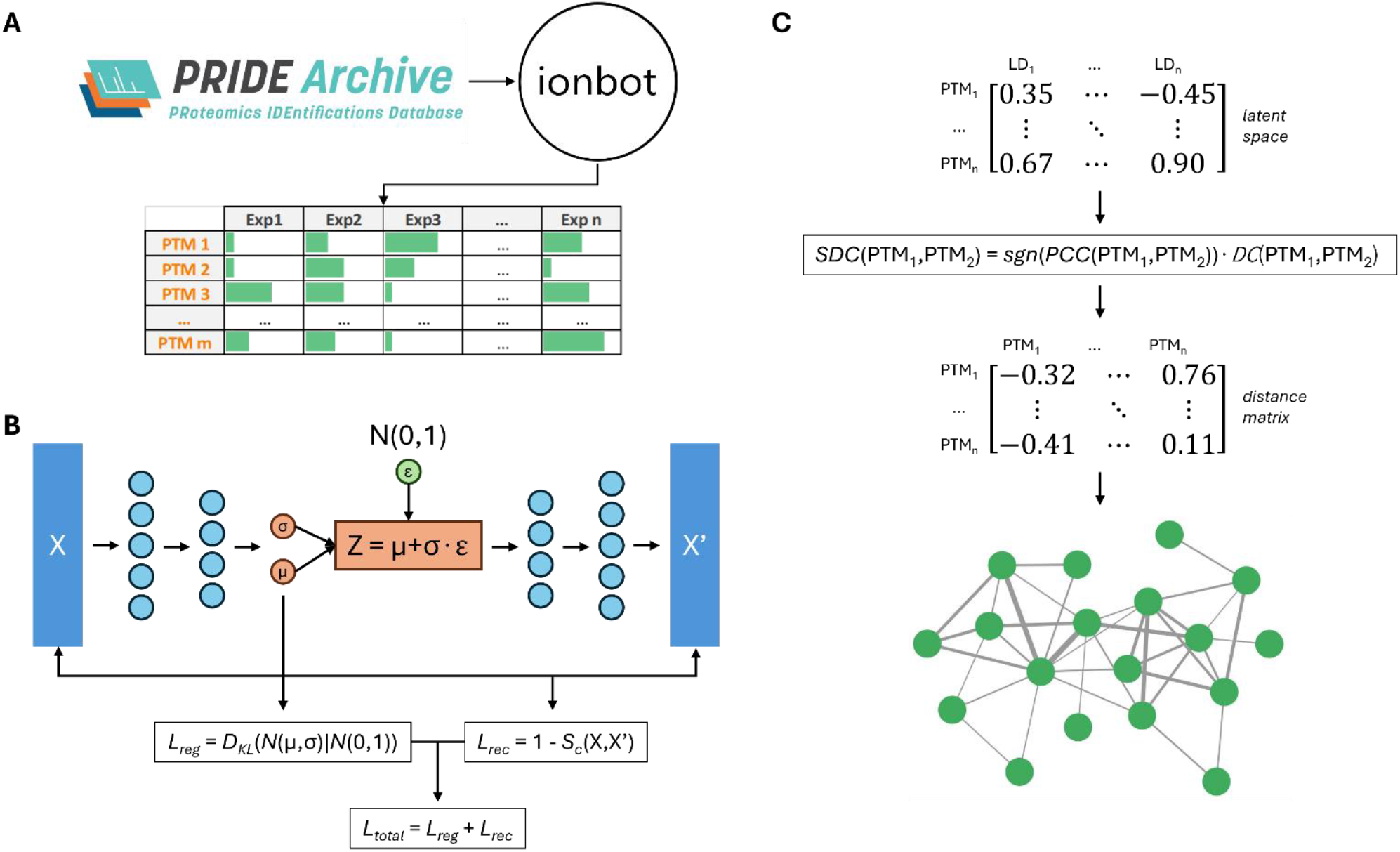
**A)** MS raw data is downloaded from the PRIDE public repository and uniformly reprocessed with ionbot to quantify PTMs across experiments. **B)** The PTM abundance matrix is used to train a VAE model, and a latent space is obtained from the VAE’s bottleneck layer. **C)** For each pair of PTMs, the signed distance correlation between the PTMs is computed. These correlations are then used as edge weights in the PTM functional associations network. (N: Normal distribution; L_reg_: regularization loss; L_rec_: reconstruction loss; L_total_: total loss; SDC: signed distance correlation; DC: distance correlation; sgn: sign function; PCC: Pearson’s correlation coefficient.)

To show that our approach can reliably identify PTM correlations, we tested it on Pulsed SILAC (pSILAC)-labelled samples [18].

pSILAC is a variation of the SILAC method where the labelled amino acids are added to the growth medium for only a short period of time, to measure differences in *de novo* protein production. Over time, light-labelled proteins are expected to decrease (due to natural turnover), while heavy-labelled ones are expected to increase.

This type of analysis thus provides a simple model in which some modifications are known to co-vary. Specifically, if we treat “unmodified” as a modification state, we expect unlabelled Arginines (R0) and Lysines (K0) to follow a similar trend; similarly, their heavy-labelled counterparts (R10, K8) will also follow a similar trend, which will be opposite to the unlabelled amino acids’ trend; however, other PTMs, such as Methionine oxidations, are not expected to correlate with either light or heavy labels.

We used three pSILAC experiments to validate our MoDPA approach (PXD021203, PXD037569, PXD042903). Raw data files were searched with ionbot specifying R10 and K8 as variable modifications rather than SILAC labels, to quantify them independently of each other. Ionbot identifications were then analysed with the MoDPA pipeline to (i) quantify PTM abundances, (ii) encode abundances in the VAE latent space, and (iii) compute signed distance correlations to infer the PTM association network.

To verify that MoDPA does not recover spurious associations, we generated a null network through degree-preserving randomization (i.e., edge rewiring that maintains the empirical degree distribution).

Validation proceeded by rule-based edge labelling: (i) positive associations between two heavy-labelled residues or two light-labelled residues were marked valid; (ii) negative associations between one heavy-labelled and one light-labelled residue were marked valid; and (iii) all other associations were marked invalid. We compared the enrichment of valid edges in the MoDPA network and in the random network and observed that the MoDPA network consistently outperforms the random one (Figure 2).

**Figure 2.**
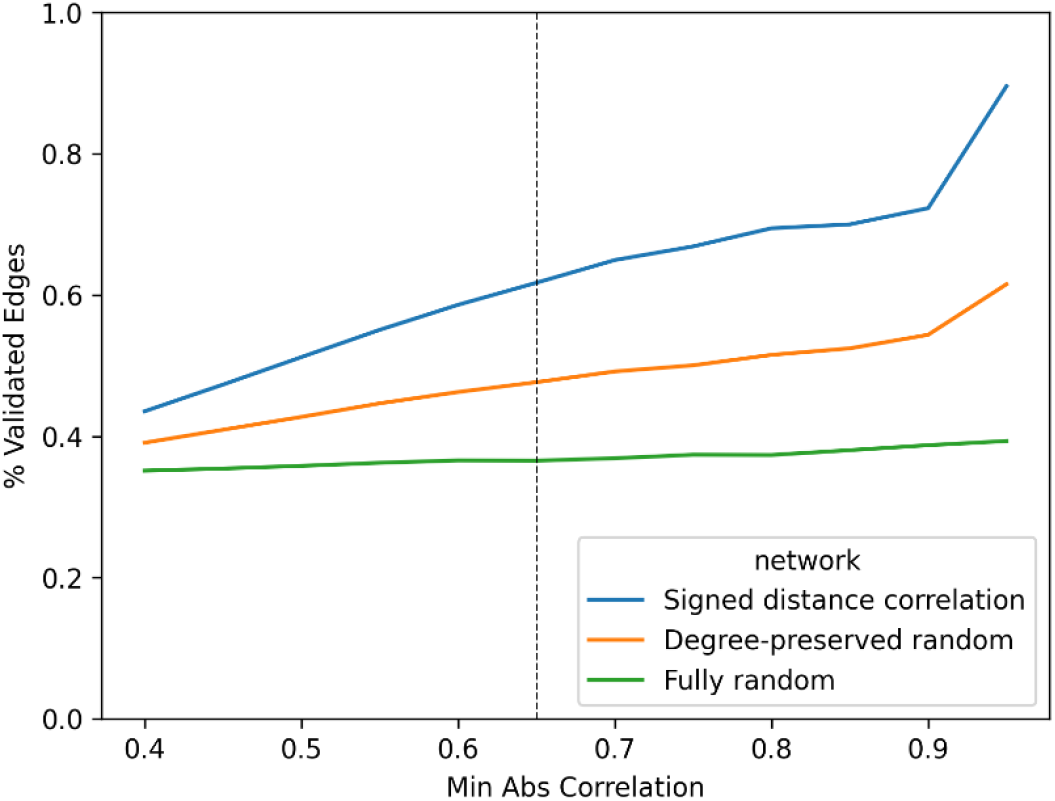
Percentage of validated edges in the network at different correlation (i.e., edge weight) thresholds. Because the network includes both positively and negatively weighted edges, we filtered on absolute weight. The dashed line indicates the threshold that was used to construct the real PTM network.

### Analysis of PTMs from PRIDE datasets

Raw data files from 640 Human PRIDE projects (Table S1) were searched with ionbot in open PTM search mode, and results were processed through the MoDPA pipeline. We recorded the occupancy of each PTM across the different MS runs, using zero-values for non-observations. We filtered for sites observed in at least 25 experiments, and for experiments with at least 25 identified PTMs.

The resulting PTM occupancy matrix was then fed to a Variational Autoencoder to obtain a latent space representation of the data. For each pair of PTMs, we calculated the signed distance correlation between their latent vectors. Associations were filtered for a correlation of at least 0.65, and for a Benjamini-Hochberg-correctedp-value smaller than 0.05.

A clustering analysis of the PTM associations network revealed the presence of 23 PTM clusters (a curated subset of clusters is shown in Figure 3A; the complete network is available in the supplementary). For each cluster, pathway over-representation analysis of the included modified proteins was performed using the human Reactome pathway knowledgebase (Figure 3B).

**Figure 3.**
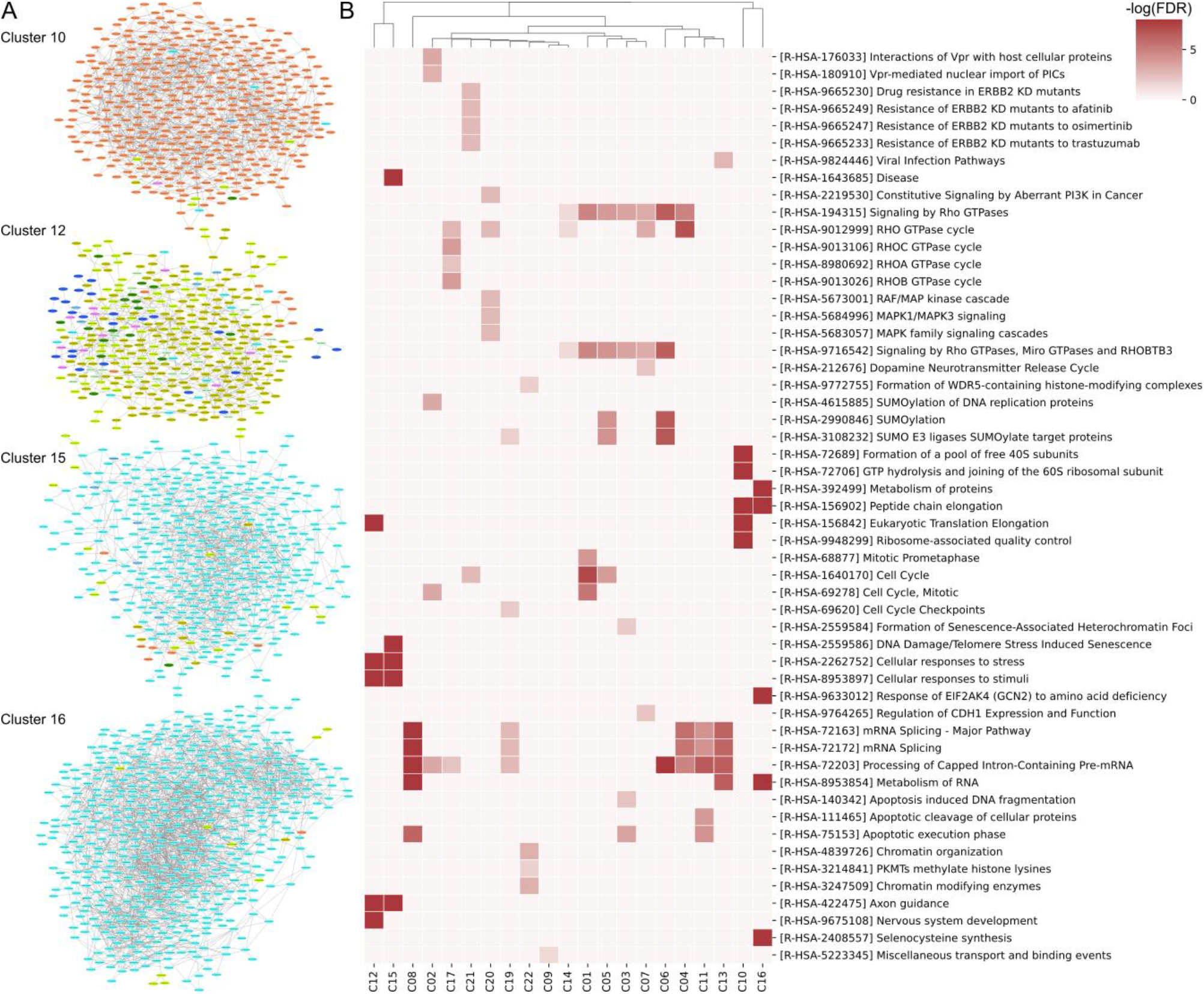
**A)** PTM clusters identified by MoDPA. The complete list of clusters is available in Supplementary Material. **B)** Reactome pathway enrichment analysis performed on the proteins in each PTM cluster. Although many pathways are enriched in multiple clusters, Clusters 10, 12, 15, and 16 are enriched in unique pathways like “Axon guidance” and “Peptide chain elongation”. These clusters also show a difference in composition (being enriched in non-phosphorylation PTMs), suggesting correlations between specific PTMs and biological processes.

Serine/threonine phosphorylation is the most abundant PTM in the network, with a total of 3838 modification sites. For this reason, most of the clusters are enriched in phosphosites. Nevertheless, four clusters stand out due to their PTM composition and the associated enriched pathways, hinting that some PTMs might be specialized in regulating some cellular processes, but not others.

**Cluster 10** is dominated by acetylation sites and shows a significant enrichment for pathways related to the metabolism of proteins and RNA (“Eukaryotic Translation Elongation”, FDR = 1.3·10^−14^; “Metabolism of RNA”, FDR = 1.3·10^−14^). While it is known that lysine acetylation is important for translation efficiency and fidelity in prokaryotes [27,28], there is limited evidence in the literature about lysine acetylation of ribosomal proteins in humans. Our identified cluster contains 27 acetylation sites on ribosomal subunits, 14 on hnRNPs (RNA-binding proteins involved in splicing, post-transcriptional regulation, etc.) [29], and 13 on translation initiation and elongation factors, suggesting the importance of this modification for mRNA translation in humans as well.

Cluster 12 is the most diverse cluster, encompassing nearly all types of the PTMs examined here, although arginine citrullination dominates, representing more than half of all sites in the cluster. The most significantly enriched terms refer to high-level pathways related to cellular stress and response to stimuli (“Cellular responses to stress” and “Cellular responses to stimuli”; FDR = 2.8·10-14), followed by modules involved in neuronal development (“Axon guidance” and “Nervous system development”; FDR = 2.8·10-14). Several other enriched pathways relate to mRNA processing and translation, including “Eukaryotic Translation Initiation” (FDR = 9.6·10-14), “Eukaryotic Translation Elongation” (FDR = 2.8·10-14), “Nonsense-Mediated Decay” (FDR = 3.0·10-12), and “Ribosome-associated quality control” (FDR = 2.2·10-12), indicating coordinated regulation of protein synthesis. Additional significant enrichments include pathways involved in selective autophagy and viral infections (“Influenza Infection”, FDR = 7.6·10-13; “SARS-CoV Infections”, FDR = 6.5·10-7), which, in Reactome, reflect host stress and immune responses that converge on translational control and proteostasis.

There is mounting evidence that protein citrullination, mediated by peptidylarginine deiminase (PAD) enzymes, plays a role in neural development processes, such as glial differentiation, myelination, and possibly gene regulation during brain development [30,31]. The cluster found here, in fact, includes known targets of PAD2 in the brain – myelin basic protein (MBP), glial fibrillary acidic protein (GFAP), and vimentin – indicating a coordinated citrullination of these proteins. Upon citrullination, arginine loses its positive charge, leading to conformational changes. In the case of myelin, this partial loss of structure provides plasticity in the developing brains of children; however, excessive arginine citrullination is linked to neurodegeneration [32]. Interestingly, citrullination is also known to be involved in the release of neutrophil extracellular traps (NETs) by neutrophils, a process dubbed NETosis, explaining the presence of immunity-related pathways in the enrichment results [33].

The pathways enriched in cluster 15 are predominantly associated with cellular stress and damage responses, with “Cellular responses to stress” (FDR = 2.1·10^−12^) and “DNA Damage/Telomere Stress Induced Senescence” (FDR = 8.5·10^−12^) among the most significantly enriched pathways. These programmes are complemented by strong enrichment of apoptotic processes, including the “Apoptotic execution phase” (FDR = 6.4·10^−10^) and “Apoptosis induced DNA fragmentation” (FDR = 8.1·10^−10^), suggesting activation of damage-induced cell fate decisions. In parallel, pathways related to host defence and vascular responses, such as “Infectious disease” (FDR = 3.2·10^−10^) and “Platelet degranulation” (FDR = 3.2·10^−9^), together with neurodevelopmental signalling captured by “Axon guidance” (FDR = 1.1·10^−10^), point to an interplay between stress-induced signalling, immune and hemostatic mechanisms, and nervous system development.

The enriched pathways in cluster 16 are primarily involved in fundamental processes of protein translation, such as “Peptide chain elongation” (FDR = 3.6·10^−15^), ribosome assembly and disassembly (“GTP hydrolysis and joining of the 60S ribosomal subunit” and “Formation of a pool of free 40S subunits”, FDR = 3.6·10^−15^), and “Nonsense-Mediated Decay” (FDR = 3.6·10^−15^). Collectively, these pathways suggest that lysine methylation plays a crucial role in the post-transcriptional regulation of gene expression.

While there is substantial evidence for the role of arginine methylation in RNA binding, splicing, and post-transcriptional regulation [34,35], evidence regarding lysine methylation is still fairly limited due to the difficulty of detecting non-histone lysine methylation events [36–38]. Despite this, our results suggest that lysine methylation might also play important roles in nucleic acid metabolism.

The remaining clusters, consisting for the most part of serine/threonine phosphorylation sites, are enriched with terms that align with this post-translational modification’s role in mRNA processing, cell cycle regulation, and signal transduction, particularly through Rho-GTPases, which govern cell morphology and movement [39].

## Discussion

VAE models have successfully been used to perform denoising and dimensionality reduction in transcriptomics and proteomics data, and to infer functional associations between proteins [16,40]. MoDPA is the first to dramatically extend this concept to protein modifications, allowing researchers to study PTM crosstalk in an unbiased and high-throughput way.

MoDPA uses a VAE to create a low-dimensional representation (latent space) of an input PTM abundance matrix. A weighted association network is then generated by calculating the SDC between each pair of PTMs in the latent space.

We applied MoDPA to two datasets: (i) a pSILAC dataset, which we used as a control due to the artificial and predictable nature of the PTMs present, and (ii) a dataset obtained from the large-scale reprocessing of 640 PRIDE projects.

In the pSILAC dataset, we demonstrate that approximately 62% of the PTM pairs with SDC ≥ 0.65 in the resulting networks are validated, i.e., they are coherent with the assumption that heavy and light labels increase and decrease over time, respectively.

We found that a cut-off of 0.65 provides a good compromise between the reliability of the associations and the connectivity of the resulting network. Nevertheless, a higher cutoff can be used to investigate specific proteins or modifications: for example, we speculate that identifying intra-protein PTM associations could pave the way for the deconvolution of proteoforms from bottom-up proteomics data. For this application, high network connectivity is not necessary, as the goal is to build one small network for each protein.

The analysis of real PTM data using MoDPA produced results that are generally consistent with the literature, while also hinting at novel lines of research, such as the role of lysine acetylation in eukaryotic translation efficiency, lysine methylation’s involvement in RNA processing, and the crosstalk between citrullination and other PTMs.

The high number of phosphosites compared to other PTMs can be attributed to the abundance of phospho-enriched datasets in PRIDE. In fact, PTMs tend to be low-abundant and require specific sample preparation steps to be successfully detected by MS. These steps are usually focused on enrichment, and are often PTM-specific (e.g., immobilized metal affinity chromatography for phosphorylation, affinity immunoprecipitation for arginine methylation), leading to the enrichment of one modification while most others are depleted [41]. This could also explain why we detect several significant associations between PTMs belonging to the same class and relatively few associations across classes (as in cluster 12). Ever-deeper coverage of proteomes as the field continues to progress will likely provide solutions for this issue over time.

Our results are released as a public resource through the online Tabloid Proteome website, to support further hypothesis generation on PTM co-regulation and disease mechanisms by researchers from the field at large.

## Conclusion

We have shown that VAE models can be used to predict PTM-dependent functional associations. Our new MoDPA method allows to correctly model the opposite abundance trends of heavy and light labels in pulsed SILAC experiments, which we used as positive controls. We then applied MoDPA to a real PTM dataset and identified clusters of significantly co-occurring modifications, many of which recapitulated known associations between PTMs and biological processes. The MoDPA PTM network is available via the Tabloid Proteome website at https://iomics.ugent.be/tabloidproteome.

## Acknowledgements

We thank all members of the CompOmics team for discussions on large-scale proteomics reprocessing and PTM site handling. We thank ELIXIR Belgium for supporting the development and maintenance of Tabloid Proteome.

## Supplementary data

**Table S1.** List of analyzed PRIDE projects.

**Table S2.** List of VAE models trained and associated hyperparameters.

**Figures S1-S23.** PTM clusters identified from the reanalysis of PRIDE data.

## Data availability

MoDPA code is available on Github (https://github.com/CompOmics/MoDPAv1.0). Data is available on Zenodo (10.5281/zenodo.18310674).

